# Purification of amylase from local isolate *Bacillus subtilis* A4

**DOI:** 10.1101/2025.08.04.668472

**Authors:** D. Jasim. M. Awda, Ali. H. Fayyadh

## Abstract

The amylase produced from local isolate *Bacillus subtilis* A4 was purified by precipitation with 80-25% saturation ammonium sulphate, followed by ion-exchange chromotography using DEAE-cellulose column, and Gel filtration using Sephdex G-100 column. In the stratification step, one peak of the enzyme was observed in the ion exchange and three major protein peaks in the recovered parts and the second summit was a container on enzymatic efficacy only the other peaks were completely empty. The number of purification times was 6 times and with an enzyme yield of 49%.

## Introduction

α-Amylase (EC 3.2.1.1, _-1,4-glucan-4-glucanohydrolase) catalyzesthe hydrolysis of α-1,4-glycosidic bonds of starch, glycogen and various related polysaccharides in a random manner and releases different sizes of oligosaccharides in an α-anomeric configuration [1]. α-Amylases represent one of the most important industrial enzymes, they have a wide variety of applications in different industrial fields, such as food, fermentation, textile, paper, detergent, pharmaceutical and sugar industries. These enzymes account for about 30% of the world’s enzyme production [2]. Although *α*-amylases can be derived from several sources, including plants, animals, and microorganisms, microbial enzymes generally meet industrial demands. e major advantage of using microorganisms for the production of *α*-amylases is the economical bulk production capacity and microbes are easy to manipulate to obtain enzymes of desired characteristics [3]. They have been reported from a wide variety of microorganisms, such as from several species of Aspergillus [4,5], from several species of genus Bacillus [2,6,7] and Streptomyces [8,9]. Up to date, microbial amylases have completely replaced chemical hydrolysis in the starch processing industry [10]. For commercial purposes α-amylases are mainly derived from the genus Bacillus [3,11]. Different species of the genus Bacillus produce α-amylases with many different properties. Some Bacillus stains produce thermo stable α-amylases, others produce_-amylases with acid-resistant property. *Bacillus subtilis* [12], *B. stearothermophilus* [13], *B. licheniformis* [14] and *B. amyloliquefaciens* [15] are known to be good producers of α-amylase with different properties, and have been widely used for commercial production of α-amylase.

## MATERIALS AND METHODS

Materials DNS (3,5-dinitrosalicylic acid), glucose, soluble starch, amylose, amylopectin, glycogen, Amicon ultrafiltration membrane and β-cyclodextrin were purchased from Sigma (Sigma-Aldrich, USA). Sephadex G-100 was purchased from Pharmacia (Pfizer and Pharmacia, Sweden). All culture medias, sodium salts and their additives were commercially obtained from Merck (Merck & Co., Inc.).

### Organism and Culture Conditions

The isolate of Bacillus subtilis used for this research was Isolated locally. The organism was grown in amedium containing (g/l): K_2_HPO_4_, 2.5; KH_2_PO_4_, 3.75; MgSO_4_, 0. 5; NaCl, 0. 5; CaCl_2_., 0.15; peptone, 10 and starch, 20. The inocula for the experiments were prepared by growing the organism in nutrient broth (NB, Oxoid) at 35Cº for 18 hrs on a rotary shaker (Gallenkamp). The medium pH was adjusted to 7.0 and autoclaved at 121 °C for 15 min and then, 50 mL medium was transferred into 500 ml flasks in a rotary shaker at 150 rpm conical flasks was inoculated with found to add the size of ainoculate containing ^8^10 x 2 cells/ml. The flask was incubated at 37oC on a rotary shaker (150 r pm) for 72hrs and then centrifuged at 20000 xg for 20 mins in cold to remove bacterial cells and the cell-free supernatant was used as a crude enzyme source. The.

### Amylase Assay

Amylase activity was estimated by the 3, 5 Dinitrosalicyclic acid (DNSA) method of (Whitaker and Bernard, 1972 ; Lin *et al*., 1997). It measures the increase in the reducing power of the digests in the reaction between starch and the enzyme. Appropriately diluted 0.1ml of enzyme was added to 0.9ml of 1% (w/v) soluble starch which was dissolved in appropriate buffer solution (phosphate buffer, 7.0). The above reaction mixture was made in three test tubes. The reaction tubes were incubated at 37Cº for 10 minutes. Then one ml of colour reagent (DNSA) was added to the reaction mixture and place in boiling water bath for 5 mins. The tubes were allowed to cool at room temperature. After which 10ml of distilled water was further added to the cooled tubes and absorbance at 540nm was measured using spectrophotometer. Control tube consisted of 0.5ml buffer solution plus 0.5ml soluble starch solution. The assay was also carried out as explained above. All assays were done in triplicates. The amount ofmaltose liberated was extrapolated from the glucose standard curve. Enzyme activity (unit/ml) it is the amount of enzyme that releases (1) micromolecules of reducing sugars(glucose) Per minute and under estimation or measurement conditions.

### Protein estimation

Protein was determined by the Biuret method of (Bradford, 1976) with bovine serum albumin (BSA) as the standard. The concentration of protein during purification studies was measured at an absorbance of 280nm.

### Purification of the amylase

The amylase enzyme purification process was studied by concentration enzyme by ammonium sulphate followed by purification by ion exchange chromatography, followed by gelatin filtration chromatography. Here is a summary of these steps:

### Ammonium sulfate precipitation

The crude culture was pre capitated with ammonium sulphate (at 80-25% saturation) to concentrate the enzyme from the raw enzymatic extract gradually in a snow bath with continuous stirring using the Stirrer engine and at 4 ° C for 30 min. Then discard the solution at 10,000 xg at 4 ° C for 20 minutes. Ignore leachate. The precipitation is the solvent of the precipitate in a small amount of (0.05 molar) of the pH solution 7.0., Then the enzymatic efficacy of all models was determined to determine the optimal saturation rate of precipitation.

### Ion-Exchange chromatography

DEAE-cellulose ionic exchange was prepared according to [15]. The enzyme obtained after Ammonium sulfate precipitation was loaded onto an equilibrated DEAE-Cellulose column (20×2.5 cm), pre-equilibrated with 0.05mM of phosphate buffer, pH 7, at room temperature, the separated parts were collected and almost run-off of 51.4ml/h. elution was performed using 0.05 molar pH phosphate precipitators of phosphate buffer, pH 7, and a linear saline gradient of sodium chloride for a 0.1-1 molar pathway, quickly with 3 ml /tube.

### Sephadex G-100 chromatography

The gel Sephdex G-100 was prepared according to the instruction of the manufacturer (Pharmacia Fine Chemical).The enzyme solution (2 ml) obtained from the step described above was loaded onto a Sephadex G-100 column (1.5×60 cm) pre-equilibrated with phosphate buffer. The column was eluted using 300 ml of the same buffer, at a flow rate of 45 ml/h. Fractions of 3.0 ml were collected, checked for enzyme activity and monitored by measuring the absorbance at 280 nm protein content.

## Results and discussion

### Extraction and purification of enzyme

The crude amylase produced by the locally isolate *Bacillus subtilis* A4 under the optimum conditions had specific activity 594.5 U/mg protein.

### Precipitation of enzyme by ammonium sulphate

In order to concentrate the crude extract of amylase and remove a much of water and some protein molecules as possible, ammonium sulphate were used at (10,15, 20, 25, 30, 40, 50, 60, 70, 80, 90)% saturation, the saturation ratio 25-80% was used. It achieved specific activity 715.5 U/mg protein, 1.2 purification fold with 88% yield. Protein.

### Ion-Exchange chromatography

Purification of amylase was done by ion-exchange chromatography by (DEAE-cellulose), Which was previously equated with the equivalent phosphate solution (0.05 molar and pH 7.0). Absorption of washable parts (un insulated proteins with positive charge) was measured along a 280 nm wavelength, when the absorption reached the base line, the recovery process of the proteins associated with the exchanger (the proteins carrying the negative charges),, The linear salt gradient recovery was performed using a phosphate solution at a concentration of 0.05 molar and pH 7.0 with linear saline gradient (0-1) molar sodium chloride. The wash step of DEAE-column contained protein peaks without amylase activity, Confirming that the enzyme was associated with the ion exchanger the step of recovery of the associated proteins was separated by protein peaks with the emergence of enzymatic activity in the recovered parts of a single peak Figure (1).

**Figure (1):**
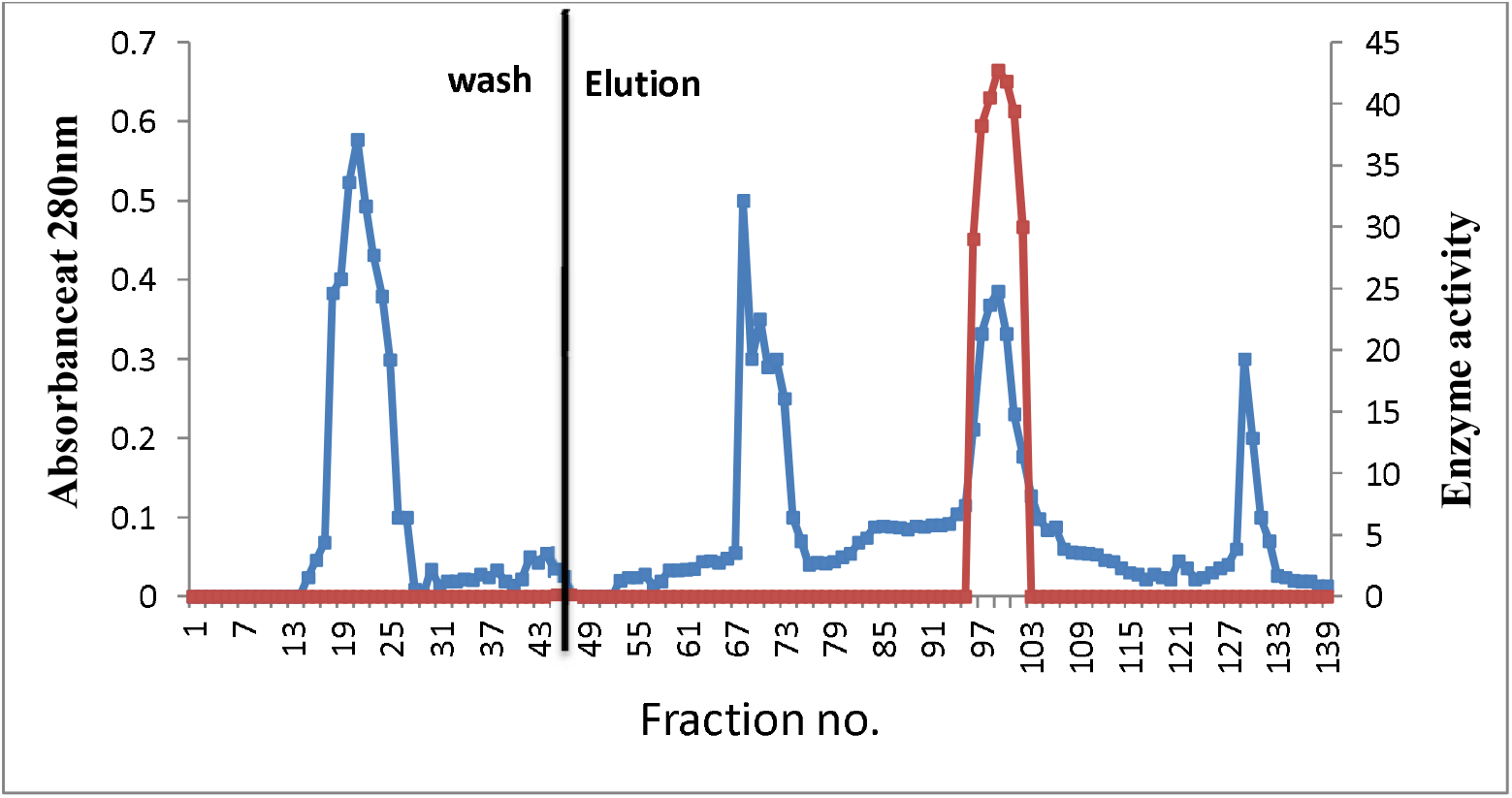
Purification of amylase from local isolate *Bacillus subtilis* A4 by DEAE-cellulose ionexchangechromatography column (20×2.5 cm) equilibrated with 0.05 M phosphate buffer pH 7, enzyme waseluted with linear salt gradient 0.1-1 M Nacl, flow rate 51.4 ml/hour

Maity *et al*., (2015) purified the enzyme amylase produced *Bacillus subtilis* (ATCC 6633) From two purification steps that included ammonium sulphate deposition and the second step using the ion exchanger DEAE cellulose The enzyme activity reached 4259 units /ml and the quality was 2662 units /mg respectively and 6 times the purification times. Singh *et al*. (2016) used the DEAE-cellulose column to purify the enzyme amylase produced from *Bacillus* sp. strain B-10 with a specific activity of 62.44 units /mg.

### Gel filtration chromatography

Transfer the enzymatic solution following the purification process of the enzyme from the previous step (ion exchanger) to the gel filtration step in the gel filter column Sephadex G-100,where the column was balanced and recovery of the enzyme with potassium potassium solution at the concentration of 0.05 molar and pH 7.0 It was noted that the parts of recovery Figure (2)) included three major protein peaks one of these peaks, the second, had only enzymatic efficacy, while the other peaks were empty Of which the peak of efficacy was largely identical to the second protein peak. Matching the efficiency curve and protein to this extent is one of the early signs of enzyme purity (Whitaker, 1972)., Collected the parts of this summit and the size and concentration of protein and enzyme activity where it was found that the size of approximately 22.5 ml Table (1).The enzyme activity and qualitative activity were estimated at 35.86 units /ml and 3586 units /mg respectively, with the number of purification times increased to 6 times and the enzyme yield at the end of this stage was 49%. Roy *et al*., (2012) indicated the purification of the enzyme amylase produced from *Bacillus subtilis* strain AS-S01a using the Sephadex G-50 column where the qualitative activity was 1500 units /mg and fold of purification 7.5 and yield 0.3%., As researcher Olufunke and Azeez, (2012) purification of the enzyme produced from *Bacillus subtilis* bacteria by the Sephadex G-150 column.The qualitative activity of 3.01 units /mg and enzymatic yield reached 42.16% with 6.40 fold.

**Figure (2):**
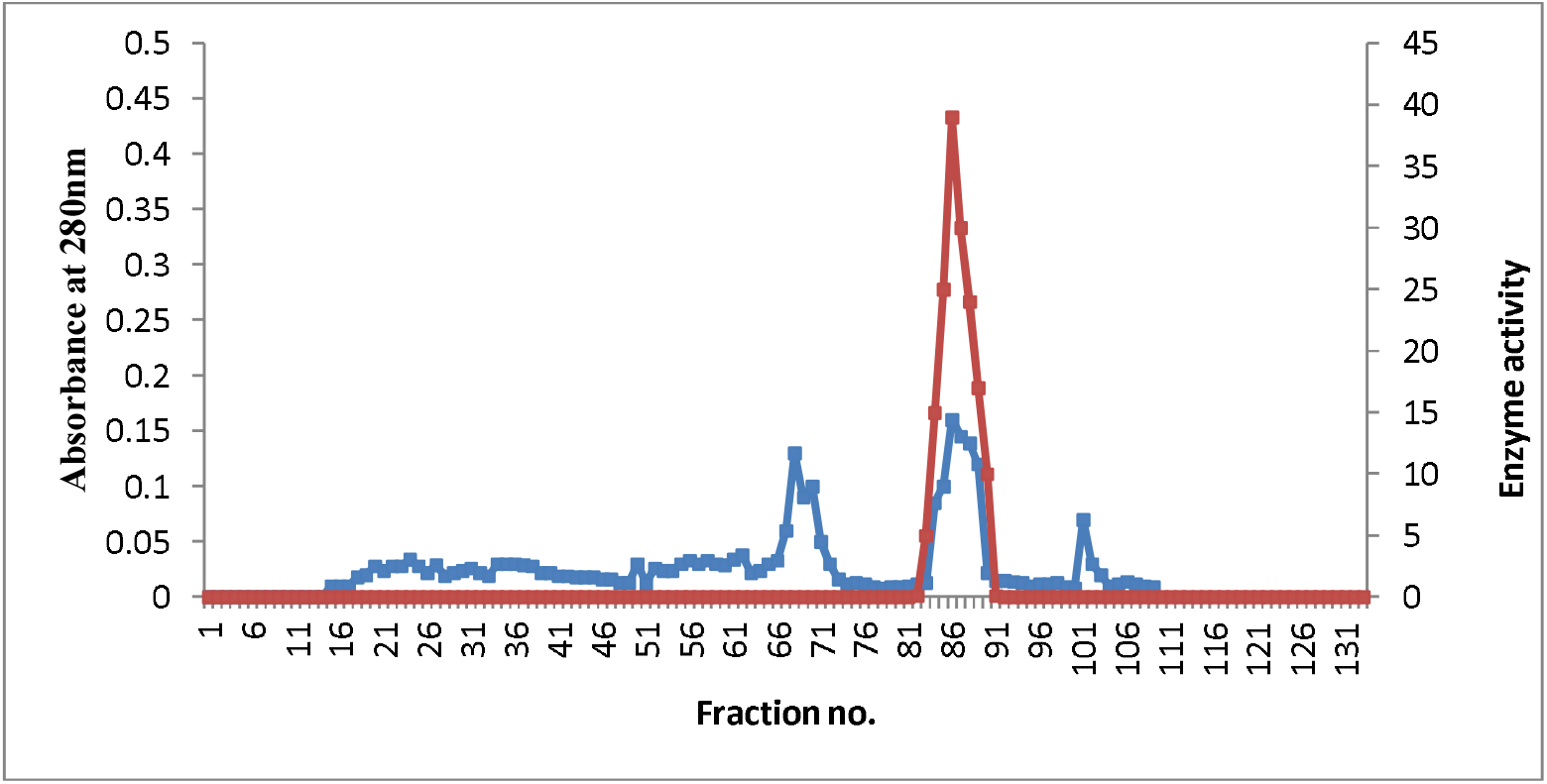
Gel filtration chromatography of amylase from local isolate *Bacillus subtilis*A4 by Sephadex G-100 column (1.5×60 cm) equilibrated with 0.05 M phosphate buffer pH 7, flow rate 45 ml/hour

**Table (1):** Purification steps of amylase produced by *Bacillus subtilis*A4.

| Steps of purification | Volume (ml) | Activity (U/ml) | Protein conc. (mg/ml) | Specific activity (U/mg) | Total activity (U) | Fold of purification | Yield (%) |
| --- | --- | --- | --- | --- | --- | --- | --- |
| Crude extract | 100 | 16.23 | 0.0273 | 594.5 | 1623 | 1 | 100 |
| Precipitation by saturation ammonium sulphate 25-80% | 20 | 71.55 | 0.1 | 715.5 | 1431 | 1.2 | 88 |
| Ion exchange chromatography by DEAE-cellulose | 27 | 41.42 | 0.02 | 2071 | 1118.34 | 3.4 | 68.9 |
| chromatography by Sephadex G-100 | 22.5 | 35.86 | 0.01 | 3586 | 806.85 | 6 | 49 |

## Notes

### Competing Interest Statement

The authors have declared no competing interest.

